# An integrated genomic analysis of anaplastic meningioma identifies prognostic molecular signatures

**DOI:** 10.1101/146811

**Authors:** Grace Collord, Patrick Tarpey, Natalja Kurbatova, Inigo Martincorena, Sebastian Moran, Manuel Castro, Tibor Nagy, Graham Bignell, Francesco Maura, Jorge Berna, Jose M. Tubio, Chris E. McMurran, Adam M.H. Young, Matthew D. Young, Imran Noorani, Stephen J Price, Colin Watts, Elke Leipnitz, Matthias Kirsch, Gabriele Schackert, Danita Pearson, Abel Devadass, Zvi Ram, V. Peter Collins, Kieren Allinson, Michael D. Jenkinson, Rasheed Zakaria, Khaja Syed, C. Oliver Hanemann, Jemma Dunn, Michael W. McDermott, Ramez W Kirollos, George S. Vassiliou, Manel Esteller, Sam Behjati, Alvis Brazma, Thomas Santarius, Ultan McDermott

## Abstract

Anaplastic meningioma is a rare and aggressive brain tumor characterised by intractable recurrences and dismal outcomes. Here, we present an integrated analysis of the whole genome, transcriptome and methylation profiles of primary and recurrent anaplastic meningioma. A key finding was the delineation of two distinct molecular subgroups that were associated with diametrically opposed survival outcomes. Relative to lower grade meningiomas, anaplastic tumors harbored frequent driver mutations in SWI/SNF complex genes, which were confined to the poor prognosis subgroup. Our analyses discern two biologically distinct variants of anaplastic meningioma with potential prognostic and therapeutic significance.

## Introduction

Meningiomas arise from arachnoidal cells of the meninges and are classified as grade I (80% of cases), grade II (10-20%) or grade III (1-3%)^1,2^. Grade III meningiomas comprise papillary, rhabdoid and anaplastic histological subtypes, with anaplastic tumors accounting for the vast majority of grade III diagnoses^2,3^. Nearly half of anaplastic meningiomas represent progression of a previously resected lower grade tumor, whereas the remainder arise *de novo*^2,4,5^. Recurrence rates are 5-20% and 20-40%, respectively, for grade I and 2 tumors^3,6^. By contrast, the majority of anaplastic meningioma patients suffer from inexorable recurrences with progressively diminishing benefit from repeated surgery and radiotherapy and 5-year overall survival of 30-60%^4,7^.

A recent study of 775 grade I and grade 2 meningiomas identified five molecular subgroups defined by driver mutation profile^8^. In keeping with previous smaller studies, mutually exclusive mutations in *NF2* and *TRAF7* were the most frequent driver events, followed by mutations affecting key mediators of PI3K and Hedgehog signalling^8,9^. Recurrent hotspot mutations were also identified in the catalytic unit of RNA polymerase II (*POLR2A*) in 6% of grade I tumors^8^. More recently, a study comparing benign versus *de novo* atypical (grade II) meningiomas found the latter to be significantly associated with *NF2* and *SMARCB1* mutations^10^. Atypical meningiomas were further defined by DNA and chromatin methylation patterns consistent with upregulated PRC2 activity, differentially methylated Homeobox domains and transcriptional dysregulation of pathways involved in proliferation and differentiation^10^.

Despite the high mortality rate of anaplastic meningiomas, efforts to identify therapeutic strategies have been hampered by a limited understanding of the molecular features of this aggressive subtype. Here, we present an analysis of the genomic, transcriptional and DNA methylation patterns defining anaplastic meningioma. Our results reveal two distinct molecular subgroups associated with dramatically different prognoses.

## Results Overview of the genomic landscape of primary and recurrent anaplastic meningioma

We performed whole genome sequencing (WGS) on a discovery set of 19 anaplastic meningiomas resected at first presentation (‘primary’). A subsequent validation cohort comprised 31 primary tumors characterised by targeted sequencing of 366 cancer genes. We integrated genomic findings with RNA sequencing and methylation array profiling in a subset of samples (Supplementary Table S1). Somatic copy number alterations and rearrangements were derived from whole genome sequencing reads, with RNA sequences providing corroborating evidence for gene fusions. Given the propensity of anaplastic meningioma to recur, we studied by whole genome sequencing 13 recurrences from 7 patients.

Excluding a hypermutated tumor (PD23359a, see Supplementary Discussion), the somatic point mutation burden of meningioma was low with a median of 28 somatic coding mutations per tumor (range 11 to 71; mean sequencing coverage 66X) in 18 tumors interrogated by WGS at first presentation (‘primary’) (Supplementary Figure S1). Mutational signatures analysis of substitutions identified in whole genome sequences revealed the age-related, ubiquitous processes 1 and 5 as the predominant source of substitutions (Supplementary Figure S2)^11^.

The rearrangement profile of anaplastic meningioma is relatively quiet, with a median of 12 structural rearrangements (range 0–79) in the 18 primary tumor genomes (Supplementary Figure S3, Supplementary Table S3). Somatic retrotransposition events, a significant source of structural variants in over half of human cancers, were scarce (Supplementary Figure S4; Supplementary Table S4)^12^. Analysis of expressed gene fusions did not reveal any recurrent events involving putative cancer genes (Supplementary Table S5).

Recurrent large copy number changes were in keeping with known copy number trends in aggressive meningiomas, notably frequent deletions affecting chromosomes 1p, 6q, 14 and 22q (Figure 1b, Supplementary Table S6)^2,8,10,13^.

**Figure 1.**
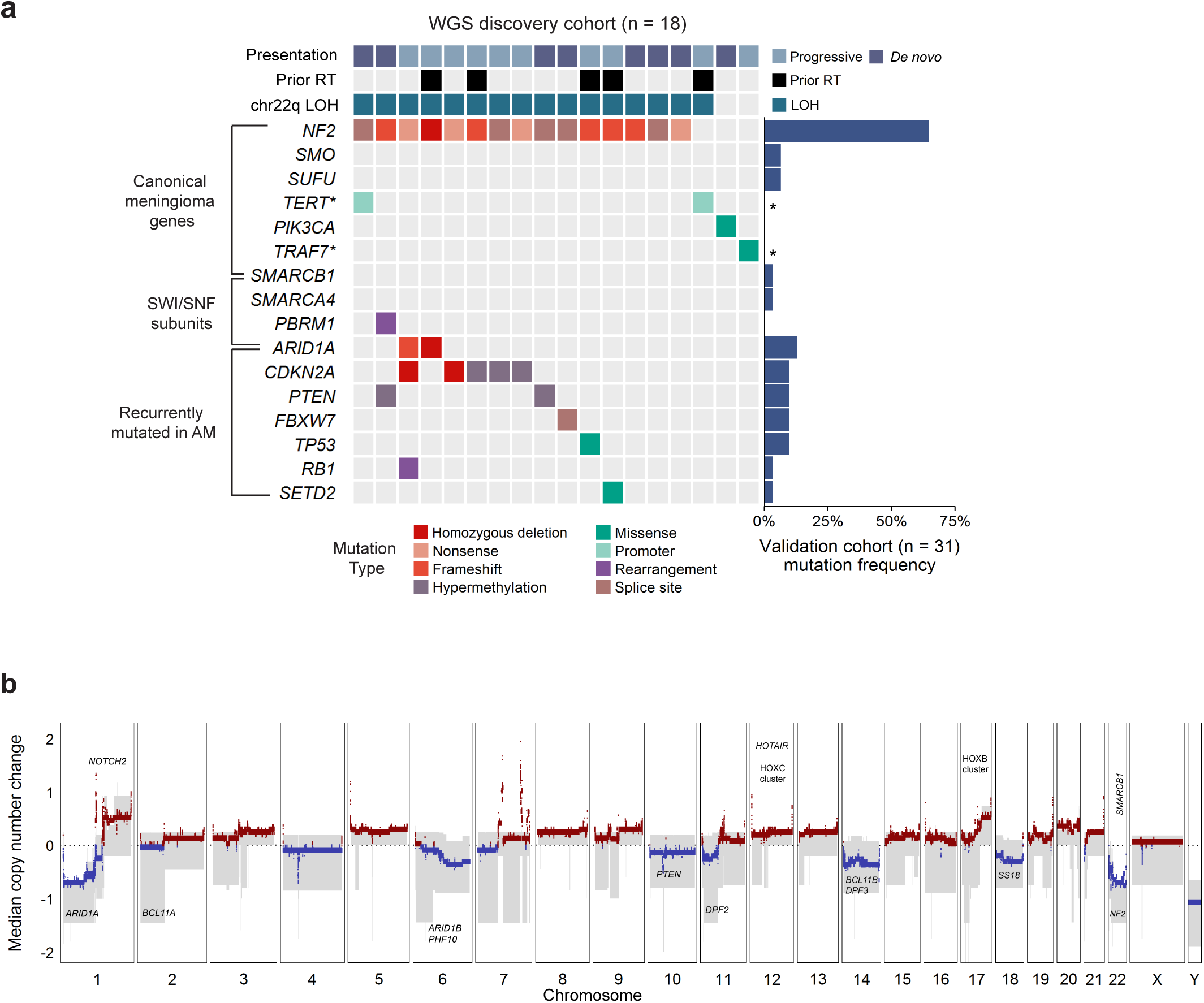
The landscape of driver mutations and copy number alterations in anaplastic meningioma. **(a)** The landscape of somatic driver variants in primary anaplastic meningioma. Somatic mutation and promoter methylation data is shown for a discovery cohort of 18 primary tumours characterised by whole genome sequenceing. Mutations in recurrently altered genes, established meningioma genes and SWI/SNF complex subunits are included. Samples are annotated for chromosome 22q LOH, prior radiotherapy exposure, and clinical presentation (*de novo* verus progression from a lower grade meningioma). The barchart to the right indicates mutation frequency in a validation cohort of 31 primary tumors sequenced with a 366 cancer gene panel. Asterisks indicate genes not included in the targeted sequencing assay. **(b)** Aggregate copy number profile of primary anaplastic meningioma. For the 18 tumors charactersied by whole genome sequencing, the median relative copy number change was calculated across the genome in 10 kilobase segments, adjusting for ploidy. The grey shaded area indicates the first and third quantile of copy number for each genomic segment. The solid red and blue lines represent the median relative copy number gain and loss, respectively, with zero indicating no copy number change. X-axis: Chromosomal position. Y-axis: median relative copy number change. Potential target genes are noted. AM, anaplastic meningioma; LOH, loss of heterozygosity; RT, radiotherapy.

The genomic landscape of recurrent tumors was largely static both with respect to driver mutations and structural variation. Driver mutations differed between primary and recurrent tumors for two of eleven patients with serial resections available. For seven sets of recurrent tumors studied by whole genome sequencing, only two demonstrated any discrepancies in large copy number variants (PD23344 and PD23346; Supplementary Figure S5). Similarly, matched primary and recurrent samples clustered closely together by PCA of transcriptome data, suggesting minimal phenotypic evolution (Supplementary Figure S6).

## Driver genes do not delineate subgroups of anaplastic meningioma

Over 80% of low grade meningiomas segregate into 5 distinct subgroups based on driver mutation profile^8,10^. In anaplastic meningioma, however, we found a more uniform driver landscape dominated by deleterious mutations in *NF2* (Figure 1a). A key feature distinguishing anaplastic meningioma from its lower grade counterparts were driver events in genes of the SWI/SNF chromatin regulatory complex (Figure 1a). The most frequently mutated SWI/SNF component was *ARID1A*, which harbored at least one deleterious somatic change in 12% of our cohort of 50 primary tumors (Supplementary Table S1). *ARID1A* has not been implicated as a driver in grade I or grade II meningiomas^8,10^. Single variants in *SMARCB1, SMARCA4* and *PBRM1* were also detected in three tumors (Supplementary Figure S7). In total, 16% of anaplastic meningiomas contained a SWI/SNF gene mutation. By contrast, SWI/SNF genes are mutated in <5% of benign and atypical meningiomas^8,10^.

In the combined cohort of 50 primary tumors, we found at least one driver mutation in *NF2* in 70%, similar to the prevalence reported in atypical meningiomas and more than twice that found in grade I tumors^8,10,14^. Interestingly, there was no significant difference in NF2 expression between *NF2* mutant and wild-type tumors (*p*-value 0.960; Supplementary Figure S8). We considered promoter hypermethylation as a source of *NF2* inactivation, but found no evidence of this (Supplementary Table S7). Thus, as observed in other cancer types, non-mutational mechanisms may contribute to *NF2* loss of function in a proportion of anaplastic meningiomas^15-20^.

Other driver genes commonly implicated in low grade tumors were not mutated, or very infrequently (Figure 1a). Furthermore, we did not observe an increased frequency of *TERT* promoter mutations, previously associated with progressive or high grade tumors^21,22^. Notably^2,23,24^, methylation analysis revealed *CDKN2A* and *PTEN* promoter hypermethylation in 17% and 11% of primary tumors, respectively (Figure 1a). We did not find evidence of novel cancer genes in our cohort, applying established methods to search for enrichment of non-synonymous mutations (See Methods and Supplementary Methods). The full driver landscape of anaplastic meningioma, considering point mutations, structural variants with resulting copy number changes and promoter hypermethylation is presented in Supplementary Figure S7.

## Differential gene expression defines anaplastic meningioma subgroups with prognostic and biological significance

We performed messenger RNA (mRNA) sequencing of 31 anaplastic meningioma samples from a total of 28 patients (26 primary tumors and 5 recurrences). Gene expression variability within the cohort did not correlate with clinical parameters including prior radiotherapy, anatomical location or clinical presentation (*de novo* versus progressive tumor) (Supplementary Figure S6). However, multiple unsupervised hierarchical clustering methods and correlation measures consistently delineated two distinct clusters, hereafter referred to as C1 and C2 (Figure 2a–c). The clinical trajectories of patients in these clusters markedly diverged. We retrospectively sought follow-up survival data from the time of first surgery, which was available for 25 of the 28 patients included in the transcriptome analysis (12 patients in C1, 13 in C2; mean follow-up of 1,403 days from surgery). We observed a significantly worse overall survival outcome in C1 compared to C2 (p<0.0001, hazard ratio 17.0 (95% CI 5.2-56.0)) (Figure 2g; Supplementary Table S8). The subgroups were well balanced with respect to potential confounding features such as gender, age, radiotherapy and anatomical location (Supplementary Table S9). Paradoxically, a greater proportion of C2 patients had higher Simpson surgical resection scores (indicating more residual tumor after surgery), conventionally a negative prognostic indicator (*P* = 0.081, Fisher’s exact test) ^4,6^.

**Figure 2.**
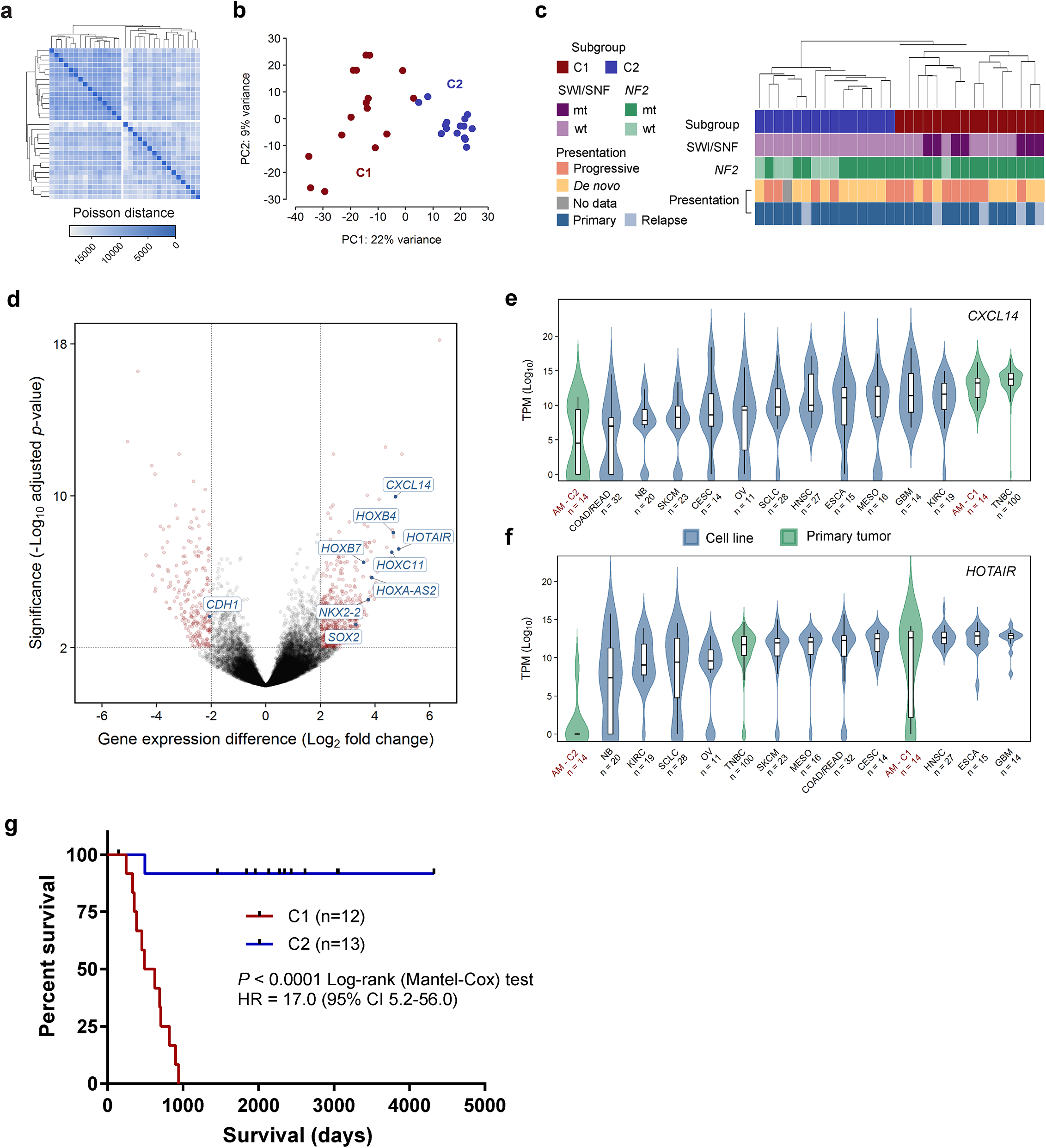
Transcriptomic classification of anaplastic meningioma. **(a)** Unsupervised hierarchical clustering and **(b)** Principal component analysis of anaplastic meningioma gene expression revealed two subgroups (denoted C1 and C2). **(c)** Dendrogram obtained by unsupervised clustering annotated with clinical and genomic features. **(d)** Volcano plot depicting genes differentially expressed between C1 versus C2 anaplastic meningioma samples. The horizontal axis shows the log_2_ fold change and the vertical axis indicates the -log_10_ adjusted *p*-value. Genes with an adjusted *p*-value < 0.01 and absolute log_2_ fold change > 2 are highlighted in red with particular genes of interest indicated. **(e, f)** Box plots of (e) CXLC14 and (f) HOTAIR expression across 31 anaplastic meningomas classified into C1 and C2 subgroups, 100 primary breast tumors, and 219 cancer cell lines from 11 tumor types. Upper and lower box hinges correspond to first and third quartiles, horizontal line and whiskers indicate the median and 1.5-fold the interquarntile range, respectively. Underlying violin plots show data distribution and are color-coded according to specimen source (blue, cell line; green, pimary tumor). X-axis indicates tumor type and number of samples in cohort. Y-axis shows NF2 Log_10_ TPM values. **(g)** Kaplan-Meier curves showing overall survival for 25 (of 28) anaplastic meningioma patients in C1 and C2 subgroups for whom follow-up data was available. Dashes indicate timepoints at which subjects were censored at time of last follow-up. TPM, transcripts per kilobase million; AM, anaplastic meningioma; TNBC, triple negative breast carcinoma; wt, wild-type; mt, mutated; HR, hazard ratio; CI, confidence interval; PC, principal component.

## Transcriptional programmes segregating anaplastic meningioma

Nineteen hundred genes underpinned the differentiation of anaplastic meningioma into subgroups C1 and C2, which could be reduced to only 6 transcripts selected on the basis of PCA coefficient and differential expression analysis (see Methods; Supplementary Tables S10 and S11; Supplementary Figure S9). Pathway enrichment analysis was most significant for evidence of epithelial-mesenchymal transition (EMT) in the C1 tumors, with concordant loss of E-cadherin (*CDH1*) and upregulation of *CXCL14*, both prognostic biomarkers in diverse other cancers (Supplementary Table S12, Figure 2d–f)^25-32^. The C1 and C2 tumors were further distinguished by significant dysregulation of proliferation, PRC2 activity and embryonic stem cell transcriptional programmes (Supplementary Table S13). Hox genes constituted a notable proportion of the transcripts distinguishing the two anaplastic meningioma subgroups, largely underpinning the significance of pathways involved in tissue morphogenesis. Furthermore, differentially methylated genes were also significantly enriched for Hox genes, with pathway analysis results corroborating the main biological themes emerging from transcriptome analysis (Supplementary Tables S14 and S15).

## Comparison of the anaplastic and benign meningioma transcriptome

Previous studies investigating the relationship between meningioma WHO grade and gene expression profiles have included few anaplastic tumors^33,34^. We therefore extended our analysis to include published RNA sequences from 19 benign grade I meningiomas. External data was processed using our in-house pipeline with additional measures taken to minimise batch effects (see Methods and Supplementary Tables S16 and S17). Unsupervised hierarchical clustering and principal component analysis demonstrated clear tumor segregation by histological grade (Figure 3a,b).

**Figure 3.**
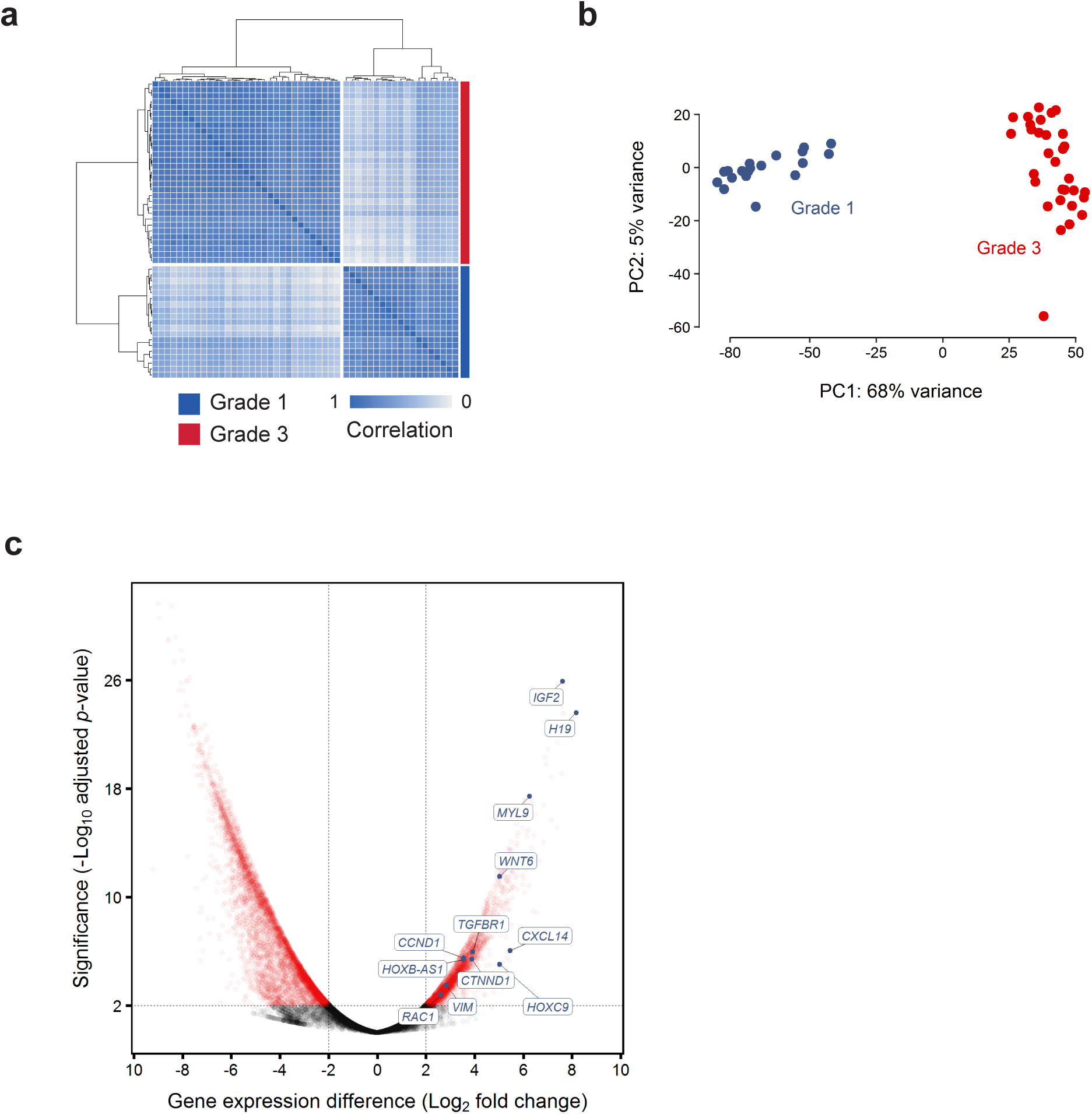
Differences in gene expression profile between grade 1 and anaplastic meningomas. **(a, b)** Normalised transcript counts from grade 1 and anaplastic meningioma samples clustered by **(a)** Pearson’s correlation coefficient and **(b)** principal component analysis. **(c)** Volcano plot illustrating differences in gene expression between anaplastic versus grade 1 meningiomas with selected genes indicated. The horizontal axis shows the log_2_ fold change and the vertical axis indicates the -log_10_ adjusted *p*-value. Genes with an adjusted *p*-value < 0.01 and absolute log_2_ fold change > 2 are highlighted in red. PC, principal component.

Consistent with there being a coherent biological trend across histological grades and anaplastic meningioma subgroups, we noted significant overlap between genes differentially expressed between grades and between C1 and C2 tumors (hypergeometric distribution *P* = 5.084E-09). In keeping with this finding, formal pathway analysis identified significant dysregulation of stemness, proliferation, EMT and PRC2 activity (Supplementary Tables S18 and S19). The most significantly dysregulated pathways also included TGF-beta, Wnt and integrin signalling, mediators of invasion and mesenchymal differentiation that are normally in part controlled by NF2 and other Hippo pathway members^15,16,35-37^. Yes-associated protein 1 (Yap1), a cornerstone of oncogenic Hippo signalling, is frequently overexpressed in cancer and synergises with Wnt signalling to induce EMT^15,37-39^. *YAP1* was upregulated in anaplastic tumors (log_2_ fold change = 2.18, FDR = 0.0062) along with *MYL9* (log_2_ fold change = 6.25, FDR = 3.76E-18), a key downstream effector essential for Yap1-mediated stromal reprogramming (Figure 3c)^38^. Additionally, the anaplastic tumors demonstrated further upregulation of major growth factor receptor and kinase circuits previously implicated in meningioma pathogenesis, notably epidermal growth factor receptor (EGFR), insulin-like growth factor (IGFR), vascular endothelial growth factor receptor (VEGFR) and mTOR complex 1 (mTORC1) kinase complex ^40-45^.

## Discussion

Meningiomas constitute a common, yet diverse tumor type with few therapeutic options^1,7,8,10^. Efforts to improve clinical outcomes have been hampered by limited understanding of the molecular determinants of aggressive disease. Here, we explored genomic, epigenetic and transcriptional features of anaplastic meningioma, the most lethal meningioma subtype^4^.

Frequent somatic changes in SWI/SNF complex genes, predominantly *ARID1A,* constitute the main genomic distinction between anaplastic and lower grade meningiomas^8,10^. SWI/SNF acts as a tumor suppressor in many cell types by antagonising the chromatin modifying Polycomb repressive complex 2 (PRC2)^46-48^. Deleterious SWI/SNF mutations unfetter PRC2 activity with profound consequences for primitive developmental pathways frequently co-opted during oncogenesis^48,49^. SWI/SNF inactivation is associated with a stem cell-like phenotype and poor outcomes in diverse cancer types^50-54^.

Although anaplastic tumors resist meaningful classification based on driver mutation patterns, unbiased transcriptional profiling revealed two biologically distinct subgroups with dramatically divergent survival outcomes. This finding is emblematic of the limitations of histopathological grading as a risk stratification system for meningioma^1,3,4,13,55^. Interestingly, all SWI/SNF mutations were exclusively identified in the poor prognosis (C1) subgroup (*P* = 0.016, Fisher’s exact test). C1 tumors were further characterised by transcriptional signatures of PRC2 target activation, stemness, proliferation and mesenchymal differentiation. These findings were in part underpinned by differential expression of Hox genes. Acquisition of invasive capacity and stem cell traits are frequently co-ordinately dysregulated in cancer, often through subversion of Hox gene programmes integral to normal tissue morphogenesis^56-58^. Hox genes have a central role in orchestrating vertebrate development and act as highly context-dependent oncogenes and tumor suppressors in cancer^57,59^. Several of the most starkly upregulated Hox genes in the C1 tumors consistently function as oncogenes across a range of solid and haematological malignancies, including *HOTAIR, HOXB7, HOXA4, HOXA-AS2, HOXC11,* and *NKX2-2*^57,60-70^. Like many long non-coding RNAs (lncRNA), *HOTAIR* and *HOXA-AS2* modulate gene expression primarily by interacting directly with chromatin remodelling complexes, exerting oncogenic activity by recruiting PRC2 to target genes^60,62,69-73^. *HOXA-AS2* has been shown to mediate transcriptional repression of the tumor suppressor gene *CDKN2A* (p16^INK4A^), loss of which is associated with poor meningioma survival^23,24,60,69,70^. Given the antagonistic relationship between the SWI/SNF and PRC2 chromatin regulators, deleterious SWI/SNF mutations and overexpression of lncRNAs known to mediate PRC2 activity emerge as potentially convergent mechanisms underpinning the differences between C1 and C2 tumors^49^.

In the context of recent studies of lower grade meningiomas, our findings raise the possibility that the balance between PRC2 and SWI/SNF activity may have broader relevance to meningioma pathogenesis. Compared to grade I tumors, atypical meningiomas are more likely to harbor *SMARCB1* mutations and large deletions encompassing chromosomes 1q, 6q and 14q. Notably, these genomic regions encompass *ARID1A* and several other SWI/SNF subunit genes. Both *SMARCB1* mutations and the aforementioned copy number changes were associated with epigenetic evidence of increased PRC2 activity, differential Homeobox domain methylation, and upregulation of proliferation and stemness programmes in the atypical tumors.

The extent to which SWI/SNF depletion plays a role in lower grade meningiomas may be therapeutically relevant. Diverse SWI/SNF mutated cancers exhibit dependence on both catalytic and non-catalytic functions of EZH2, a core subunit of PRC2^74-76^. Several EZH2 inhibitors are in development with promising initial clinical results^77^. Other modulators of PRC2 activity, including HOTAIR, may also be relevant therapeutic targets^78,79^. Furthermore, growing recognition of the relationship between EMT and resistance to conventional and targeted anti-cancer agents has profound implications for rational integration of treatment approaches^80,81^. Notably, EGFR inhibition has yielded disappointing response rates in meningioma^41,82^. A mesenchymal phenotype is strongly associated with resistance to EGFR inhibitors in lung and colorectal cancer^80,81,83-85^. Combining agents that abrogate EMT with other therapies is a promising strategy for addressing cell-autonomous and extrinsic determinants of disease progression and may warrant further investigation in meningioma^80,86^.

This study has revealed prognostically significant anaplastic meningioma subgroups and identified potentially actionable alternations in SWI/SNF genes and other therapeutically tractable targets. However, a substantially larger series of tumors, ideally nested in a prospective multicentre observational study, will be required to expand upon our main findings and explore mechanistic and therapeutic ramifications of meningioma diversity.

## Methods Sample selection

DNA was extracted from 70 anaplastic meningiomas; 51 samples at first resection (‘primary’) and 19 from subsequent recurrences. Matched normal DNA was derived from peripheral blood lymphocytes. Written informed consent was obtained for sample collection and DNA sequencing from all patients in accordance with the Declaration of Helsinki and protocols approved by the NREC/Health Research Authority (REC reference 14/YH/0101) and Ethics Committee at University Hospital Carl Gustav Carus, Technische Universität Dresden, Germany (EK 323122008). Samples underwent independent specialist pathology review (V.P.C and K.A). DNA extracted from fresh-frozen material was submitted for whole genome sequencing whereas that derived from formalin-fixed paraffin-embedded (FFPE) material underwent deep targeted sequencing of 366 cancer genes.

One tumor sample PD23348 (and two subsequent recurrences) separated from the main study samples in a principal components analysis of transcriptomic data (Supplementary Figure S10). Analysis of WGS and RNA sequencing data identified an expressed gene fusion, *NAB2-STAT6*. This fusion is pathognomonic of meningeal hemangiopericytoma, now classified as a separate entity, solitary fibrous tumors^87-89^. We therefore excluded three samples from this tumor from further study. A second sample (PD23354a), diagnosed as an anaplastic meningioma with papillary features, was found to have a strong APOBEC mutational signature as well as an *EML4-ALK* gene fusion (exon 6 EML4, exon 19 ALK) (Supplementary Figure S11)^90^. Therefore this sample was also removed as a likely metastasis from a primary lung adenocarcinoma. The hypermutator sample PD23359a underwent additional pathological review to confirm the diagnosis of anaplastic meningioma (K.A., Department of Histopathology, Cambridge University Hospital, Cambridge, UK).

RNA was extracted from fresh-frozen material from 34 primary and recurrent tumors, 3 of which were from PD23348 and were subsequently excluded from final analyses (Supplementary Table S1).

### Whole genome sequencing

Short insert 500bp genomic libraries were constructed, flowcells prepared and sequencing clusters generated according to Illumina library protocols^91^. 108 base/100 base (genomic), or 75 base (transcriptomic) paired-end sequencing were performed on Illumina X10 genome analyzers in accordance with the Illumina Genome Analyzer operating manual. The average sequence coverage was 65.8X for tumor samples and 33.8X for matched normal samples (Supplementary Table S1).

## Targeted genomic sequencing

For targeted sequencing we used a custom cRNA bait set (Agilent) to enrich for all coding exons of 366 cancer genes (Supplementary Table S20). Short insert libraries (150bp) were prepared and sequenced on the Illumina HiSeq 2000 using 75 base paired-end sequencing as per Illumina protocol. The average sequence coverage was 469X for the tumor samples.

## RNA sequencing and data processing

For transcriptome sequencing, 350bp poly-A selected RNA libraries were prepared on the Agilent Bravo platform using the Stranded mRNA library prep kit from KAPA Biosystems. Processing steps were unchanged from those specified in the KAPA manual except for use of an in-house indexing set. Reads were mapped to the GRCh37 reference genome using STAR (v2.5.0c)^92^. Mean sequence coverage was 128X. Read counts per gene, based on the union of all exons from all possible transcripts, were then extracted BAM files using HTseq (v0.6.1)^93^. Transcripts Per kilobase per Million reads (TPM) were generated using an in-house python script (https://github.com/TravisCG/SI_scripts/blob/master/tpm.py)^92,93^. We downloaded archived RNA sequencing FASTQ files for 19 grade I meningioma samples representing the major mutational groups (*NF2*/chr22 loss, *POLR2A, KLF4/TRAF7, PI3K* mutant) (ArrayExpress: GSE85133)^8^. Reads were then processed using STAR and HTseq as described above. Cancer cell line (n=252) and triple-negative breast cancer (n = 100) RNA sequencing data was generated in-house by the aforementioned sequencing and bioinformatic pipeline.

Expressed gene fusions were sought using an in-house pipeline incorporating three algorithms: TopHat-Fusion (v2.1.0), STAR-Fusion (v0.1.1) and deFuse (v0.7.0) (https://github.com/cancerit/cgpRna)^92,94,95^. Fusions identified by one or two algorithms or also detected in the matched normal sample were flagged as likely artefacts. Fusions were further annotated according to whether they involved a kinase or known oncogene and whether they occurred near known fragile sites or rearrangement break points^96^ (Supplementary Table S5).

## Differential gene expression and pathway enrichment analysis

The DESeq2 R package was used for all differential gene expression analyses^97,98^. DESeq2 uses shrinkage estimation of dispersion for the sample-specific count normalization and subsequently applies a linear regression method to identify differentially expressed genes (DEGs)^97,98^.

Preliminary comparison of anaplastic and externally-generated grade I meningioma data revealed evidence of laboratory batch effects, which we mitigated with two batch-correction methods: RUVg and PEER^99,100^. RUVg estimates the factor attributed to spurious variation using control genes that are assumed to have constant expression across samples^101-103^. We selected control genes (*RPL37A, EIF2B1, CASC3, IPO8, MRPL19, PGK1* and *POP4*) on the basis of previous studies of suitable control genes for transcript-based assays in meningioma^104^. PEER (‘probabilistic estimation of expression residuals’) is based on factor analysis methods that infer broad variance components in the measurements. PEER can find hidden factors that are orthogonal to the known covariates. We applied this feature of PEER to remove additional hidden effect biases. The final fitted linear regression model consists of the factor identified by RUVg method that represents the unwanted laboratory batch effect and 13 additional hidden factors found by PEER that are orthogonal to the estimated laboratory batch effect. Using this approach we were able to reduce the number of DEGs from more than 18000 to 8930, of which <4,000 are predicted to be protein-coding.

To identify biological pathways differentially expressed between meningioma grades and anaplastic meningioma subgroups we applied a functional class scoring algorithm using a collection of 461 published gene sets mapped to 10 canonical cancer hallmarks (Supplementary Table S21)^56,105-109^. We further corroborated these findings with a more general Gene Ontology (GO) pathway analysis^110^.

## Identification of 6 transcripts recapitulating anaplastic meningioma clusters

Mapped RNA sequencing reads were normalised using the regularised logarithm (rlog) function implemented by the DESeq2 package^97,98^. PCA was performed using the top 500 most variably expressed transcripts and the prcomp function (R stats package)^111^. Given that primary component 1 (PC1) was the vector most clearly distinguishing the closely clustered C2 subgroup from the more diffusely clustered C1 (Figure 3a), we extracted the top 50 transcripts with the highest absolute PC1 coefficients. We then identified the subset that overlapped with the most significantly differentially expressed genes (absolute log_2_ fold change > 4 and adjust *p*-value < 0.0001) between i) the C1 and C2 anaplastic meningioma subgroups and ii) the C1 anaplastic meningiomas and the 19 grade I tumors (Supplementary Tables S10 and S17). Iteratively reducing the number of PC1 components identified the minimum number of transcripts that recapitulated segregation of C1 and C2 tumors upon unsupervised hierarchical clustering and PCA (Supplementary Table S11; Supplementary Figure S9).

## Processing of genomic sequencing data

Genomic reads were aligned to the reference human genome (GRCh37) using the Burrows-Wheeler Aligner, BWA (v0.5.9)^112^. CaVEMan (Cancer Variants Through Expectation Maximization: http://cancerit.github.io/CaVEMan/) was used for calling somatic substitutions. Small insertions and deletions (indels) in tumor and normal reads were called using a modified Pindel version 2.0. (http://cancerit.github.io/cgpPindel/) on the NCBI37 genome build^113,114^. Annotation was according to ENSEMBL version 58. Structural variants were called using a bespoke algorithm, BRASS (BReakpoint AnalySiS) (https://github.com/cancerit/BRASS) as previously described^115^.

The ascatNGS algorithm was used to estimate tumor purity and ploidy and to construct copy number profiles from whole genome data^116^.

## Identification of cancer genes based on the impact of coding mutations

To identify recurrently mutated driver genes, we applied a dN/dS method that considers the mutation spectrum, the sequence of each gene, the impact of coding substitutions (synonymous, missense, nonsense, splice site) and the variation of the mutation rate across genes^115^. To detect genes under significant selective pressure by either point mutations or indels, each gene’s *P*-values from dN/dS analysis of substitutions and from the recurrence analysis of indels were combined using Fisher’s method. Multiple testing correction (Benjamini-Hochberg FDR) was performed separately for all genes, stratifying the FDR correction to increase sensitivity^117^. To achieve a low false discovery rate a conservative *q*-value cutoff of <0.05 was used for significance and considered significant any gene with qmis_sfdr<0.05 OR qglobal_sfdr<0.05. See Supplementary Methods.

## Identification of driver mutations in known cancer genes

Non-synonymous coding variants detected by Caveman and Pindel algorithms were flagged as putative driver mutations according to set criteria and further curated following manual inspection in the Jbrowse genome browser^118^. Variants were screened against lists of somatic mutations identified by a recent study of 11,119 human tumors encompassing 41 cancer types and also against a database of validated somatic drivers identified in cancer sequencing studies at the Wellcome Trust Sanger Institute (Supplementary Tables S22 and S23)^119^.

Copy number data was analysed for homozygous deletions encompassing tumor suppressor genes and for oncogene amplifications exceeding 5 or 9 copies for diploid and tetraploid genomes, respectively. Only focal (<1Mb) copy number variants meeting these criteria were considered potential drivers. Additional truncating events (disruptive rearrangement break points, nonsense point mutations, essential splice site mutations and out of frame indels) in established tumor suppressors were also flagged as potential drivers. Only rearrangements with breakpoints able to be reassembled at base pair resolution are included in this dataset.

## TraFiC pipeline for retrotransposon integration detection

For the identification of putative solo-L1 and L1-transduction integration sites, we used the TraFiC (Transposome Finder in Cancer) algorithm^12^. TraFiC uses paired-end sequencing data for the detection of somatic insertions of transposable elements (TEs) and exogenous viruses. The identification of somatic TEs (solo-L1, Alu, SINE, and ERV) is performed in three steps: (i) selection of candidate reads, (ii) transposable element masking, (iii) clustering and prediction of TE integration sites and (iv) filtering of germline events^12^.

## Methylation arrays and analysis

We performed quantitative methylation analysis of 850,000 CpG sites in 25 anaplastic meningiomas. Bisulfite-converted DNA (bs-DNA) was hybridized on the Ilumina Infinium HumanMethylationEPIC BeadChip array following the manufacturer’s instructions. All patient DNA samples were assessed for integrity, quantity and purity by electrophoresis in a 1.3% agarose gel, picogreen quantification and Nanodrop measurements. Bisulfite conversion of 500 ng of genomic DNA was done using the EZ DNA Methylation Kit (Zymo Research), following the manufacturer’s instructions. Resulting raw intensity data (IDATs) were normalized under R statistical environment using the Illumina normalization method developed under the minfi package (v1.19.10). Normalized intensities were then used to calculate DNA methylation levels (beta values). Then, we excluded from the analysis the positions with background signal levels in methylated and unmethylated channels (p>0.01). Finally we removed probes with one or more single nucleotide polymorphisms (SNPs) with a minor allele frequency (MAF) > 1% in the first 10 bp of the interrogated CpG, as well as the probes related to X and Y chromosomes. From the filtered positions, we selected only CpG sites present both in promoter regions (TSS, 5’UTR and 1st exon) and CpG islands (UCSC database, genome version hg19).

For the supervised analysis of the probes, CpG sites were selected by applying an ANOVA test to identify statistically significant CpG positions (FDR adjusted p-value < 0.01) that were differentially methylated among the compared groups (Δβ > 0.2). Selected CpG sites were later clustered based on the Manhattan distances aggregated by ward’s linkage. Finally, the genes corresponding to the selected CpGs were used to perform a Gene Set Enrichment Analysis (GSEA) with curated gene sets in the Molecular Signatures Database^120^. The gene sets used were: H: hallmark gene sets, BP: GO biological process, CC: GO cellular component, MF: GO molecular function and C3: motif gene sets (http://software.broadinstitute.org/gsea/msigdb/collections.jsp). The gene clusters resulting from the hypergeometric test with a FDR adjusted p-value < 0.05 were finally considered. We observed high levels of methylation for *CREBBP* in the majority of tumor samples, however, similar patterns were manifest in normal tissue controls, hence *CREBBP* hypermethyation does not appear to be a feature of oncogenesis in these samples.

## Mutational signature analysis

Mutational signature extraction was performed using the nonnegative matrix factorization (NNMF) algorithm^11^. Briefly, the algorithm identifies a minimal set of mutational signatures that optimally explains the proportions of mutation types found across a given mutational catalogue and then estimates the contribution of each identified signature to the mutation spectra of each sample.

## Patient survival analysis

The Kaplan-Meier method was used to analyze survival outcomes by the log-rank Mantel-Cox test, with hazard ratio and two-sided 95% confidence intervals calculated using the Mantel_Haenszel test (GraphPad Prism, ver 7.02). Overall survival data from time of first surgery for each anaplastic meningioma within gene-expression defined subgroups 1 and 2 was collected and used to plot a Kaplan-Meier survival curve.

## Data availability

All sequencing data that support the findings of this study have been deposited in the European Genome-Phenome Archive and are accessible through the accession numbers EGAS00001000377, EGAS00001000828, EGAS00001000859, EGAS00001001155 and EGAS00001001873. All other relevant data are available from the corresponding author on request.

## Supplementary Discussion

### A hypermutator anaplastic meningioma with a haploid genome

One primary anaplastic meningioma resected from an 85-year old female (PD23359a) had a hypermutator phenotype, with 27,332 point mutations and LOH across nearly its entire genome (Supplementary Figure S12; Supplementary Table S24). Independent pathological review confirmed the original diagnosis of anaplastic meningioma, and transcriptome analysis confirmed that this tumor clustered appropriately with the rest of the cohort (Figure 3a,b). The majority of the mutations were substitutions, 72% of which were C>T transitions. We identified two deleterious mutations in DNA damage repair mediators: a *TP53* p.R248Q missense mutation and a homozygous truncating variant in the mismatch repair gene *MSH6* (p.L1330Vfs*9). Despite the latter finding, mutational signatures analysis was dominated by signature 1, with no evidence of signatures typically associated with defects in homologous recombination, mismatch repair or *POLE* activity (signatures 3, 6, 10, 15, 20 or 26). The copy number profile is most consistent with this tumor having first undergone haploidization of its genome, with the exception of chromosomes 7, 19 and 20, followed by whole genome duplication (Supplementary Figure S12). Of note, several important oncogenes are located on chromosome 7, including *EGFR, MET* and *BRAF*. Widespread LOH has been described in a significant proportion of oncocytic follicular thyroid cancers where preservation of chromosome 7 heterozygosity has also been observed^121^.

## Acknowledgements

This work was supported by the Wellcome Trust, Cancer Research UK, Meningioma UK and Tadhg and Marie-Louise Flood. U.M. was personally supported by a Cancer Research UK Clinician Scientist Fellowship; G.C. by a Wellcome Trust Clinical PhD Fellowship (WT098051); F.M. by A.I.L. (Associazione Italiana Contro le Leucemie-Linfomi e Mieloma ONLUS) and by S.I.E.S. (Società Italiana di Ematologia Sperimentale); S.B. was funded by a Wellcome Trust Intermediate Clinical Research Fellowship and a St. Baldrick’s Foundation Robert J. Arceci Innovation Award. J.B. was funded by the charity Brain Tumour Research. The samples were received from the tissue banks from Cambridge (UK), Dresden (Germany), Liverpool (UK), Plymouth (UK) and Tel Aviv (Israel). The Human Research Tissue Bank is supported by the NIHR Cambridge Biomedical Research Centre. We are grateful to the patients who enabled this study and to the clinical teams coordinating their care.

## Author Contributions

G.C. and N.K. performed mRNA expression analysis. P.T. and G.C analysed whole genome and targeted sequencing data. I.M. performed statistical analyses to detect novel driver mutations. S.M. analysed methylation array data. F.M. generated mutational signatures analysis. J.T. and M.C. performed retrotransposon analysis. C.O.H. and J.D. performed protein expression analysis. A.B, S.B. and M.Y. contributed to data analysis strategy. A.Y., T.N., G.R.B and J.T. provided informatic support. T.S., R.W.K., M.K, G.S., D.P., A.D., C.E.M., A.Y., I.N., S.J.P., C.W., Z.R., M.D.J., R.Z., and K. S. provided samples and clinical data. S.B., G.S.V, I.N. and M.W.M. provided conceptual advice. V.P.C and K.A carried out central pathology review. U.M. and T.S. devised and supervised the project. U.M. G.C. and T.S. wrote the manuscript with input from S.B., P.T., and G.S.V. All authors approved the manuscript.

## Competing Financial Interests

All authors declare no competing financial interests.

Correspondence and requests for materials should be addressed to U.M. and T.S.

